# Proteomic Profiling Identifies CLDN3 as a Tumor-Selective Therapeutic Target in Small Cell Lung Cancer

**DOI:** 10.64898/2026.07.29.738916

**Authors:** Brett A. Schroeder, Jae Choi, Alejandro A. Schäffer, Michael Nirula, Yingying Cao, Yang Zhang, Anna-Lena Meinhardt, Hyeokjun Kwon, Hyerim Park, Sangin Lee, Hyo Jin Shin, Hee Geon Park, Hobin Yang, Sungyoul Hong, Parth Desai, Donna Butcher, Pattara Sukprasert, Binbin Wang, D. Nathan Biery, Rajaa El Meskini, Devon Atkinson, Laura L. Bassel, Zoe Weaver Ohler, Yue Huang, Allison L. Hunt, Tamara Abulez, Nicholas W. Bateman, Thomas P. Conrads, Ross Lake, Stephen M. Hewitt, Sunghyun Hong, Jaemun Kim, Seung Hwa Jung, Ho Jin Lee, Saehyung Lee, Young Kee Shin, Eytan Ruppin, Anish Thomas

## Abstract

Small cell lung cancer (SCLC) remains a highly lethal disease with limited targetable surface antigens beyond delta-like ligand 3 (DLL3). We sought to systematically identify and validate tumor-selective cell-surface targets in relapsed SCLC. To this end, we developed an integrated proteogenomic pipeline combining single-cell RNA sequencing, combinatorial optimization, mass spectrometry–based proteomics, and immunohistochemistry to systematically map the SCLC surfaceome of 49 tumors across 25 patients with relapsed SCLC. Using this approach we found claudin-3 (*CLDN3*) to be a consistently expressed and tumor-selective antigen, with broader coverage than DLL3, which is the current clinical benchmark. CLDN3 was highly expressed across various treatment states, while maintaining low expression in most nonmalignant tissues. Functional validation using a novel CLDN3-specific monoclonal antibody (ABN501) demonstrated NK cell–mediated cytotoxicity *in vitro* and tumor regression *in vivo*, as well as a favorable safety profile. These findings support clinical development of CLDN3-directed therapies and demonstrate the utility of integrated proteogenomic approaches for antigen discovery in solid tumors.

## INTRODUCTION

Small cell lung cancer (SCLC) is the most lethal form of lung cancer, with a median survival of less than one year and more than 30,000 new cases diagnosed annually in the United States^1,2^. Although most patients initially respond to platinum-based chemotherapy, relapse is nearly universal and durable remissions are rare. Immune checkpoint inhibitors (ICIs) are approved in the treatment of SCLC, but clinical benefit remains limited, with most patients deriving limited long-term benefit^3–6^.

The therapeutic landscape shifted with the FDA approval of tarlatamab, a bispecific T-cell engager (TCE) that recruits CD8^+^ T cells to DLL3-expressing tumor cells^7,8^. DLL3, an inhibitory Notch ligand that sustains the SCLC neuroendocrine state by blocking Notch signaling, is aberrantly expressed on SCLC cells while being largely absent from normal tissues^9^. DLL3 was nominated based on mechanistic insights into SCLC biology. However, despite its strong biological rationale, tarlatamab response rates remain below 40%, highlighting persistent challenges related to intratumoral heterogeneity^10^, antigen variability, and incomplete target coverage^11^. These limitations underscore the need for additional and complementary targets.

TCEs are part of a broader class of targeted modalities, including antibody-drug conjugates (ADCs), monoclonal antibodies, nanoparticles, CAR T-cells, and targeted protein degraders^12–16^. These approaches engage targets via distinct mechanisms: receptor-mediated endocytosis, direct cell surface engagement, or intracellular protein degradation. However, many of these strategies depend, directly or indirectly, on tumor-selective cell-surface antigens. Therefore, identifying robust surface targets remains a crucial challenge.

Yet, the SCLC surfaceome remains incompletely defined. Systematic mapping efforts have long been limited by several obstacles: tumors are rarely surgically resected, are not represented in large-scale efforts such as The Cancer Genome Atlas (TCGA), and biopsies at relapse are infrequent due to rapid disease progression. As a result, detailed molecular characterization of SCLC has been limited, hindering systematic discovery of targets. Moreover, target expression can vary at relapse^17^, and individual tumors often contain heterogeneous malignant cell states^2^. Thus, a single target may fail to cover all clinically relevant tumor populations, particularly in relapsed disease. These features underscore the need to identify additional surface antigens that are retained across treatment states and can complement existing targets such as DLL3.

Here, we applied an integrative approach that combined single-cell transcriptomics, surfaceome prediction, and proteomic validation to systematically map the relapsed SCLC surface landscape (**Fig. S1**). This multi-layered analysis was designed to reveal tumor-selective surface antigens, including those associated with therapy-resistant cell populations, and to establish a foundation for the development of next-generation targeted therapies. We identify claudin-3 (CLDN3) as a tumor-selective, broadly expressed surface antigen and provide a rational basis for the development of CLDN3-directed therapies, such as ABN501, a new fully human monoclonal antibody.

## RESULTS

### Study Cohort and Identification of Malignant Cell Populations

We collected single-cell RNA sequencing (scRNA-seq) data from 49 tumor samples from 25 SCLC patients spanning diverse metastatic sites, including liver, lymph node, adrenal, lung, mediastinum, breast, pleura, and kidney. All patients had received platinum-based chemotherapy, and 23/25 (92%) had been treated with immunotherapy (**Table S1**). After filtering and quality control (see **Methods**), 39 samples were retained for downstream surface target prediction.

Unsupervised UMAP clustering of the scRNA-seq data revealed 16 transcriptionally distinct populations comprising tumor, immune, and stromal cells (**Fig. S2A-B**). Copy number variation (CNV) analysis demonstrated characteristic chromosomal gains and losses, distinguishing tumor from non-tumor populations (**Fig. S2C-D**). Tumor cell identity was further supported by expression canonical SCLC transcripts such as *MYC*, *INSM1*, and *ASCL1,* which were enriched within malignant clusters (**Fig. S2E**).

### Combinatorial Optimization Identifies Patient-Specific Surface Targets and Nominates CLDN3

We applied MadHitter, a combinatorial optimization algorithm, to single-cell transcriptomic data from relapsed SCLC patients to identify candidate surface antigens and estimate target complexity for each tumor (**Fig. 1A**)^18^. Specifically, MadHitter models target selection as a hitting set problem, where it determines the smallest set of targets needed to “hit” (i.e., cover) a required fraction of tumor cells while sparing non-tumor cells. We used standard thresholds requiring coverage of at least 80% of malignant cells while targeting no more than 10% of non-malignant cells. Additional parameters included a minimum tumor-to-non-tumor expression ratio of 2.0. Analyses were restricted to 2,752 surface protein-encoding genes after filtering^19^, and each sample was evaluated independently to identify targets that could meet tumor-selectivity and safety thresholds.

**Figure 1.**
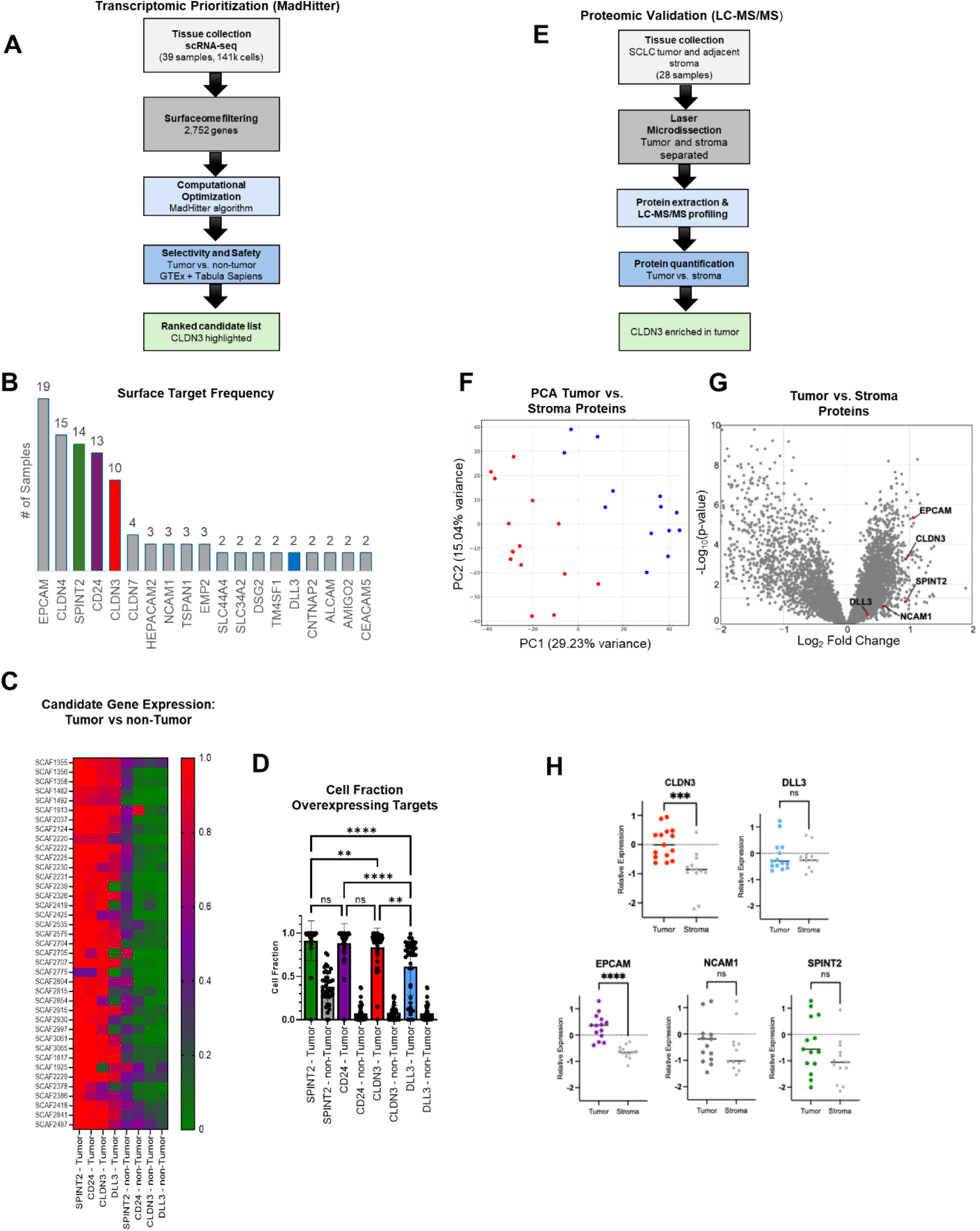
Integrated transcriptomic and proteomic discovery identifies CLDN3 as a tumor-selective therapeutic target in SCLC. (A) Overview of the MadHitter pipeline: scRNA-seq data (39 SCLC samples) filtered for 2,752 surfaceome genes, ranked by tumor selectivity, and screened for normal-tissue safety (GTEx, Tabula Sapiens). (B) Frequency of top 7 target transcripts across samples; CLDN3 was most recurrent, with CD24, SPINT2, and DLL3 also enriched. (C) Heatmap of candidate transcript expression (normalized 0–1) shows tumor-selective enrichment of CLDN3 and DLL3, while SPINT2 is broadly expressed in non-tumor cells. (D) Cell fractions with OE ≥ 0.5; CLDN3, DLL3, and CD24 were significantly enriched in tumors versus non-tumor (two-way ANOVA), while SPINT2 was non-selective. (E) Schematic overview of the experimental workflow. Tumor and adjacent stromal regions from small cell lung cancer (SCLC) samples (n = 28) were harvested by laser microdissection (LMD for protein digestion and LC-MS/MS-based proteomic profiling to quantify different protein abundance. (F) Principal component analysis (PCA) of all quantified proteins demonstrates clear separation between tumor (red) and stroma (blue) samples, confirming effective compartmental enrichment. (G) Volcano plot of differential protein expression comparing tumor and stromal samples. Key surface targets including CLDN3, DLL3, EPCAM, SPINT2, and NCAM1 are enriched in tumor tissue. (H) Paired analysis of protein expression in tumor versus stroma for individual targets: CLDN3, DLL3, EPCAM, NCAM1, and SPINT2. Paired two-sided t-tests were used. Asterisks denote significance: ns = not significant; p < 0.05 (*), p < 0.01 (**), p < 0.001 (***), p < 0.0001 (****).

Across the cohort, 31 of 39 samples (79%) were targetable with a single antigen under standard thresholds (≥80% tumor killing, ≤10% normal impact). Under more stringent thresholds (≥90%/≤5%), 18 of 39 samples (46%) remained targetable by a single antigen. Because multiple distinct antigen sets can achieve the same optimal solution for a given sample, we ranked the top seven candidate genes per sample, identifying 21 recurrently prioritized genes (**Fig. 1B**). *EPCAM* was most frequently selected target (19 of 39 samples), followed by *CLDN4* (15 samples), *SPINT2* (14), *CD24* (13), and *CLDN3* (10). By contrast, *DLL3* was prioritized in only two samples, consistent with more limited predicted coverage in relapsed SCLC.

Despite its frequent selection, *EPCAM* was deprioritized due to its expression across normal epithelial tissues, raising concerns for on-target/off-tumor toxicity^20^. CLDN4 also emerged as a strong candidate based on its prevalence and predicted tumor specificity; however, prior antibody-based efforts targeting CLDN4 have shown only moderate preclinical activity, with limited clinical translation to date^21,22^. SPINT2, a serine protease inhibitor, has been independently identified as a potential SCLC surface target through detection of circulating autoantibodies bound to native membrane antigens from patient plasma^23^. The B-cell differentiation marker CD24 is also highly expressed in SCLC^24^, but prior therapeutic efforts using radiolabeled anti-CD24 antibodies showed early accumulation in bone marrow and spleen, raising concerns about off-tumor toxicity^25^. Together, these data nominated CLDN3 as a compelling candidate for further evaluation, prompting subsequent analyses that compared its expression and therapeutic potential with DLL3 and included SPINT2 and CD24 as additional comparators.

To refine candidate selection, we compared transcript expression patterns of these four genes across tumor and non-tumor compartments (**Fig. 1C**). *CLDN3* and *CD24* demonstrated high expression in malignant cells of many samples, with minimal expression in non-tumor populations, supporting strong tumor selectivity. *DLL3*, though enriched in tumors, showed lower overall abundance and more heterogeneity across samples. *SPINT2* displayed higher non-tumor expression, raising concerns for off-tumor on-target toxicity. Quantitative analysis confirmed that *CLDN3* and *CD24* had significantly greater tumoral expression relative to healthy tissue, compared to *DLL3* and *SPINT2* (**Fig. 1D**).

### *CLDN3* is More Broadly Expressed Than *DLL3* And Expressed on Poorly Differentiated Cell States

To evaluate target coverage and safety, we compared *CLDN3* and *DLL3* across SCLC samples using threshold criteria that balance tumor selectivity with sparing of normal tissue. To make these predictions clinically interpretable, we applied threshold criteria to estimate how well a target balances tumor coverage with sparing of normal tissue. The “80/10” threshold requires that a candidate antigen be expressed on at least 80% of tumor cells while appearing on no more than 10% of non-tumor cells. At this standard level, *CLDN3* as a single target met these criteria in 13 samples (33%) compared to only 7 samples (18%) for *DLL3* (**Fig. S3A**). A stricter “90/5” threshold requires ≥90% tumor coverage and ≤5% non-tumor expression, reflecting a higher bar for both efficacy and safety. Even under these stringent conditions, *CLDN3* remained a viable target in 9 samples (23%), whereas *DLL3* met criteria in only 1 sample (3%). On average, *CLDN3* achieved broader tumor coverage compared to *DLL3*, and showed low expression in non-tumor cells (**Fig. S3B**), underscoring it promising therapeutic potential, combining high coverage of tumor cells (potential for effective killing) with low coverage of non-tumor cells (potential safety).

We next asked whether *CLDN3* expression was associated with aggressive tumor cell states. CytoTRACE^26^ analysis revealed that high *CLDN3* expression was enriched in tumor clusters with high stemness scores (**Fig. S3C-E**), consistent with poorly differentiated, therapy-resistant compartments thought to fuel relapse. UMAP visualization of the CLDN3:DLL3 ratio highlighted regions with relative *CLDN3* dominance (**Fig. S3F**), and violin plots confirmed heterogeneity of this ratio across malignant clusters (**Fig. S3G**). Thus, CLDN3 provides broader tumor coverage than DLL3 and may also preferentially mark less differentiated SCLC cell states. Together, these findings nominate CLDN3 as a tumor-selective antigen with superior coverage and therapeutic potential compared to DLL3.

### Mass Spectrometry Confirms Tumor-Specific Expression of CLDN3

To validate surface targets identified by MadHitter, we performed MS-based proteomic profiling on laser microdissection (LMD) enriched tumor and stromal compartments from 11 relapsed SCLC patients (28 total samples) (**Fig. 1E**). Tissue preparation, protein extraction, and LC-MS/MS analysis were carried out as previously described^27^. In total, 7,418 proteins were quantified in tumor and 6,655 in stromal samples, with 6,155 proteins co-quantified in both compartments. Principal component analysis (PCA) demonstrated clear separation between tumor and stromal proteomes, confirming effective compartmental enrichment (**Fig. 1F**).

Differential protein expression analysis^28^ revealed that several candidate antigens were preferentially enriched in tumor tissue compared to stroma (**Fig. 1G**). CLDN3 showed the strongest and most consistent tumor-specific enrichment (p < 0.0001). DLL3 was also elevated in tumors (p = 0.0188) but displayed greater variability across samples. EPCAM also showed significant enrichment in tumor (p < 0.01). In contrast, NCAM1 and SPINT2 were not consistently different between compartments (**Fig. 1H**).

These proteomic data provide orthogonal evidence that corroborates the transcriptomic predictions from MadHitter. Taken together, the convergence of scRNA-seq and mass spectrometry identifies CLDN3 as a robust, tumor-selective surface antigen that outperforms DLL3 in breadth and consistency of expression.

### CLDN3 Shows Tumor Selectivity Across Treatment States in an Independent SCLC Cohort

To validate our findings in an independent cohort, we analyzed the expression of *CLDN3, DLL3, CD24*, and *SPINT2* in 18 scRNA-seq samples from SCLC patients from an independent cohort^29^, categorized into treatment-naïve, post-treatment, and metastatic groups (**Fig. S4**). We assessed the proportion of tumor and non-tumor cells overexpressing each candidate target using MadHitter (see Methods).

Consistent with our discovery cohort, *CLDN3* was robustly enriched in tumor cells across all treatment conditions, with minimal expression in non-tumor populations. *DLL3*, while tumor-associated, again showed lower overall abundance and greater variability. These findings support our earlier conclusion that *CLDN3* provides broader and more consistent tumor coverage than *DLL3*. They also suggest that a combinatorial approach incorporating both *CLDN3* and *DLL3* could further expand tumor coverage, particularly in metastatic disease, although this requires future experimental validation.

*CD24* and *SPINT2* were included as comparators, and although both showed high tumor cell expression in some cases, *SPINT2* exhibited substantial non-tumor expression, especially in post-treatment and metastatic samples. *CD24*, while tumor-enriched, showed variable non-tumor expression across clinical states. These findings support CLDN3 as a highly tumor-selective and broadly expressed surface target in SCLC, independent of treatment status, and support its prioritization for therapeutic development.

### Assessment of Target Safety Using Protein and Transcript Expression in Normal Tissues

Next, we evaluated CLDN3 expression across normal tissues to assess potential off-tumor toxicity, using complementary approaches. Immunohistochemistry (IHC) of representative healthy organs, including brain, adrenal, kidney, liver, lung, and lymph node, demonstrated that CLDN3 was absent or weakly expressed in most tissues, while strong membranous staining was observed in SCLC and metastatic lesions (**Fig. 2A**). Semiquantitative scoring across a panel of normal organs confirmed that the majority of tissues showed “Not Detected” or “Low” expression, although higher expression was noted in gastrointestinal and parathyroid samples, consistent with the physiologic localization of claudins in epithelial tight junctions (**Fig. 2B**).

**Figure 2.**
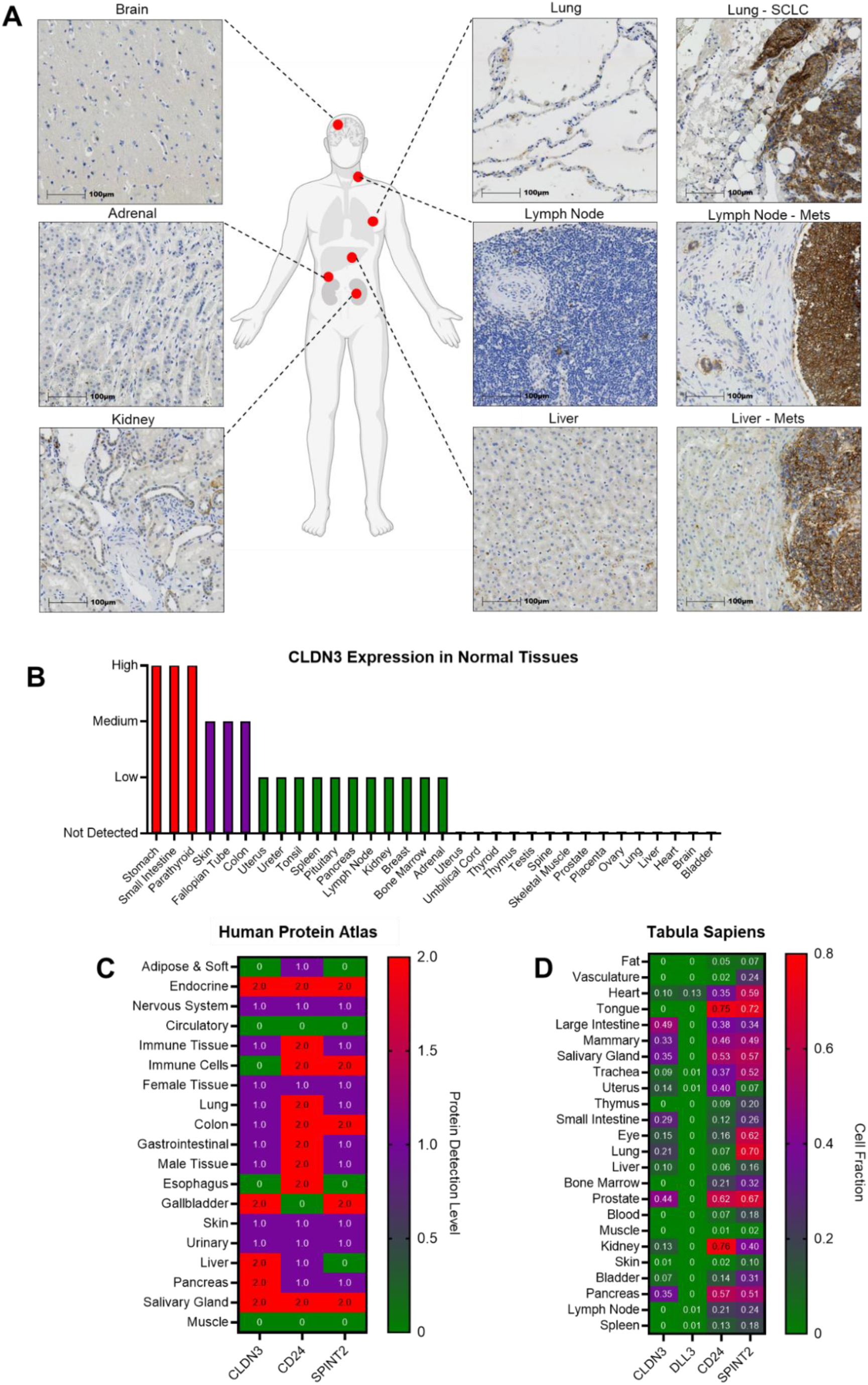
CLDN3 shows limited expression in normal tissue. (A) Representative IHC of CLDN3 protein expression in normal tissues (brain, lung, adrenal, kidney, liver, lymph node) versus SCLC and metastatic lesions. CLDN3 was undetectable or weak in most normal tissues but strongly expressed in tumor samples with membranous staining. (B) Semiquantitative summary of CLDN3 protein expression across normal tissues (Not Detected: 0%; Low: <25%; Medium: 25–75%; High: >75%). Most tissues fell into Not Detected or Low, with few tissues scoring High. (C) Heatmap of RNA expression across tissue categories (Human Protein Atlas). CLDN3, CD24, and SPINT2 showed low expression across most tissues. DLL3 not available. (D) Detection frequencies of CLDN3, DLL3, CD24, and SPINT2 in specific normal tissue subtypes (Tabula Sapiens atlas). DLL3 showed minimal or absent expression across healthy tissues.

Analysis of public transcriptomic datasets further supported this profile. Data from the Human Protein Atlas^30^ showed that CLDN3, along with CD24 and SPINT2, exhibited low protein expression across most tissue categories, with minimal detection in critical compartments such as nervous system, immune cells, and muscle (**Fig. 2C**). Tabula Sapiens scRNA-seq similarly revealed low transcript detection frequencies of CLDN3 across healthy tissue (**Fig. 2D**). Specifically, CLDN3 expression was negligible in brain and heart, despite restricted epithelial expression in gastrointestinal and parathyroid tissues. These observations are consistent with independent proteomic studies reporting limited CLDN3 expression in normal tissues and its tumor specificity, consistent with its suitability as a therapeutic target^31^.

Together, these orthogonal datasets indicate that CLDN3 is selectively enriched in tumors with minimal expression in normal tissues. Although low-level proteomic expression is detectable in gastrointestinal and parathyroid epithelia, its absence from critical parenchymal organs, combined with strong tumor-specific expression, supports CLDN3 as a promising and potentially safe therapeutic target in SCLC.

### IHC Validation of MadHitter Predictions Confirms Superior Tumor Coverage by CLDN3

Given the clinical relevance of DLL3 as a validated SCLC target, we performed IHC to directly compare CLDN3 and DLL3 expression at the protein level and validate MadHitter predictions. A total of 13 tumors from 10 patients were analyzed, including 9 for which DLL3 was predicted to have high expression and all 13 with CLDN3 predictions. Representative IHC images from four tumors demonstrated strong and widespread membranous staining of CLDN3, often exceeding that of DLL3 within the same samples (**Fig. 3A**).

**Figure 3.**
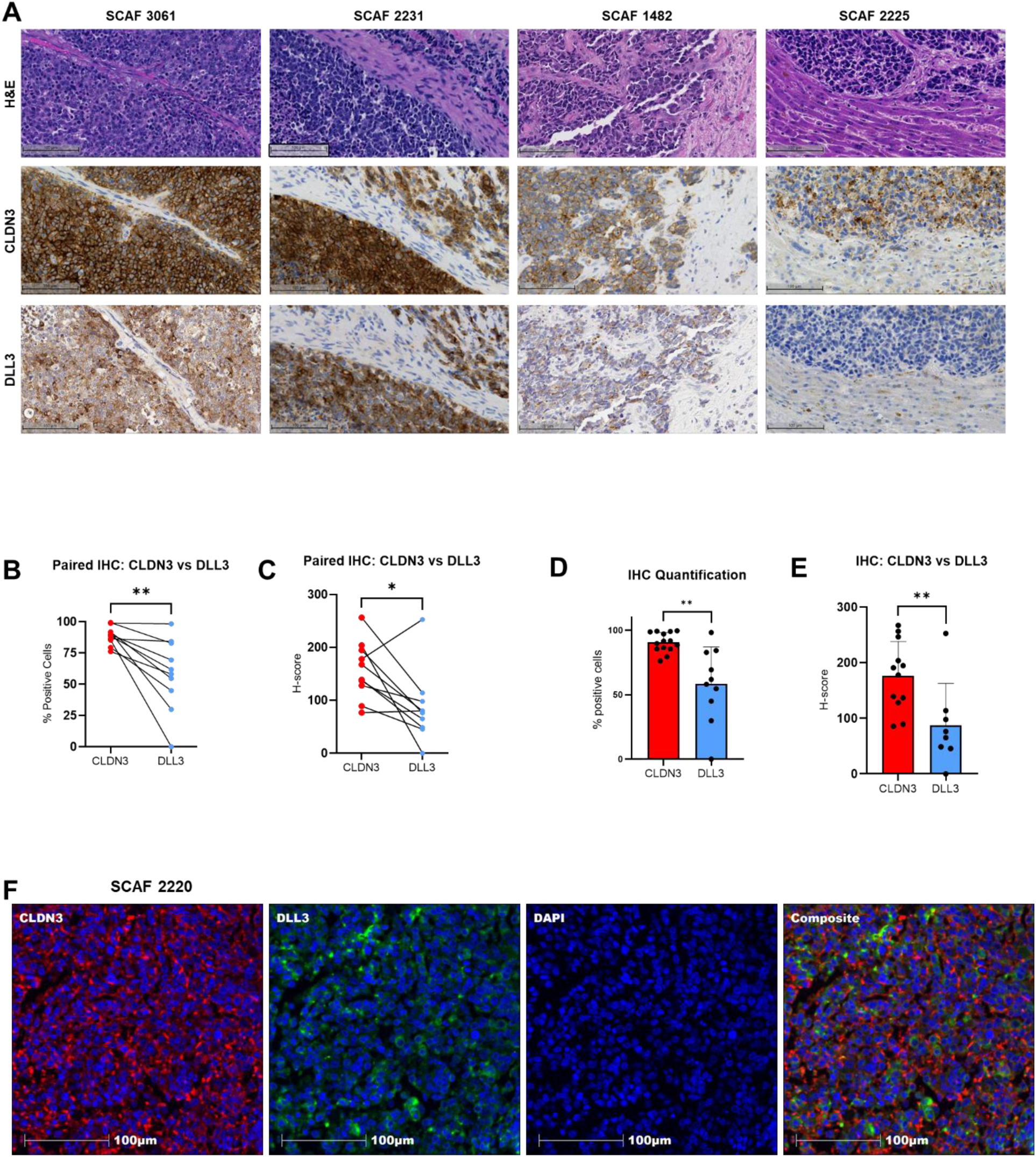
Immunohistochemistry of CLDN3 vs. DLL3. (A) Representative IHC from four SCLC tumors (SCAF3061, 2231, 1482, 2225), showing H&E (top), DLL3 (middle), and CLDN3 (bottom). Scale bar = 100 µm. (B) Paired comparison of % DLL3^+^ vs. CLDN3^+^ cells with matched scRNA-seq predictions (n = 6); paired t-test, p < 0.01. (C) Paired comparison of DLL3 vs. CLDN3 H-scores in matched samples with MadHitter predictions; paired t-test. (D) % positive cells (left) and H-scores (right) for CLDN3 vs. DLL3 across all IHC samples (n = 11); unpaired t-tests, p < 0.01 and p < 0.05. (E) H-scores in samples with scRNA-seq predictions (n = 13); unpaired t-test, p < 0.05. (F) Immunofluorescence of tumor SCAF2220 showing DAPI (blue), DLL3 (green), CLDN3 (red), and composite. Scale bar = 100 µm.

In paired analyses of samples with predictions for both genes, CLDN3 was expressed in a significantly higher fraction of tumor cells compared to DLL3 (paired t-test, p < 0.01; **Fig. 3B**). H-scores, which integrate staining intensity and distribution, were likewise higher for CLDN3 across paired tumors (**Fig. 3C**). When all available samples (n=11) were included, CLDN3 again demonstrated significantly greater coverage than DLL3, both by percent positivity and H-score (unpaired t-tests, p < 0.01 and p < 0.05, respectively; Fig. **3D-E**).

Immunofluorescence imaging further confirmed these findings, showing robust and widespread CLDN3 protein expression relative to DLL3 in the same tumor (**Fig. 3F**). Collectively, these data validate MadHitter predictions and highlight CLDN3 as a consistently expressed, tumor-selective surface target that is highly expressed in more tumors than DLL3, on both the transcriptomics and proteomic levels.

### CLDN3 Surface Expression Predicts Susceptibility to NK Cell–Mediated Cytotoxicity

As a membrane protein, the therapeutic potential of CLDN3 depends not only on selective expression but also on whether antibodies can engage it on the cell surface and mediate immune effector killing. To test this, we performed antibody-dependent cellular cytotoxicity (ADCC) assays across a panel of 16 SCLC cell lines representing all four molecular subtypes, ASCL1 (A), NEUROD1 (N), POU2F3 (P), and YAP1 (Y), capturing the phenotypic diversity observed in patient tumors^32^. Flow cytometry showed a wide range of CLDN3 surface expression across lines and general concordance with mRNA abundance reported in DepMap^33^, supporting the alignment of transcriptomic predictions with protein-level expression (**Fig. 4A**).

**Figure 4.**
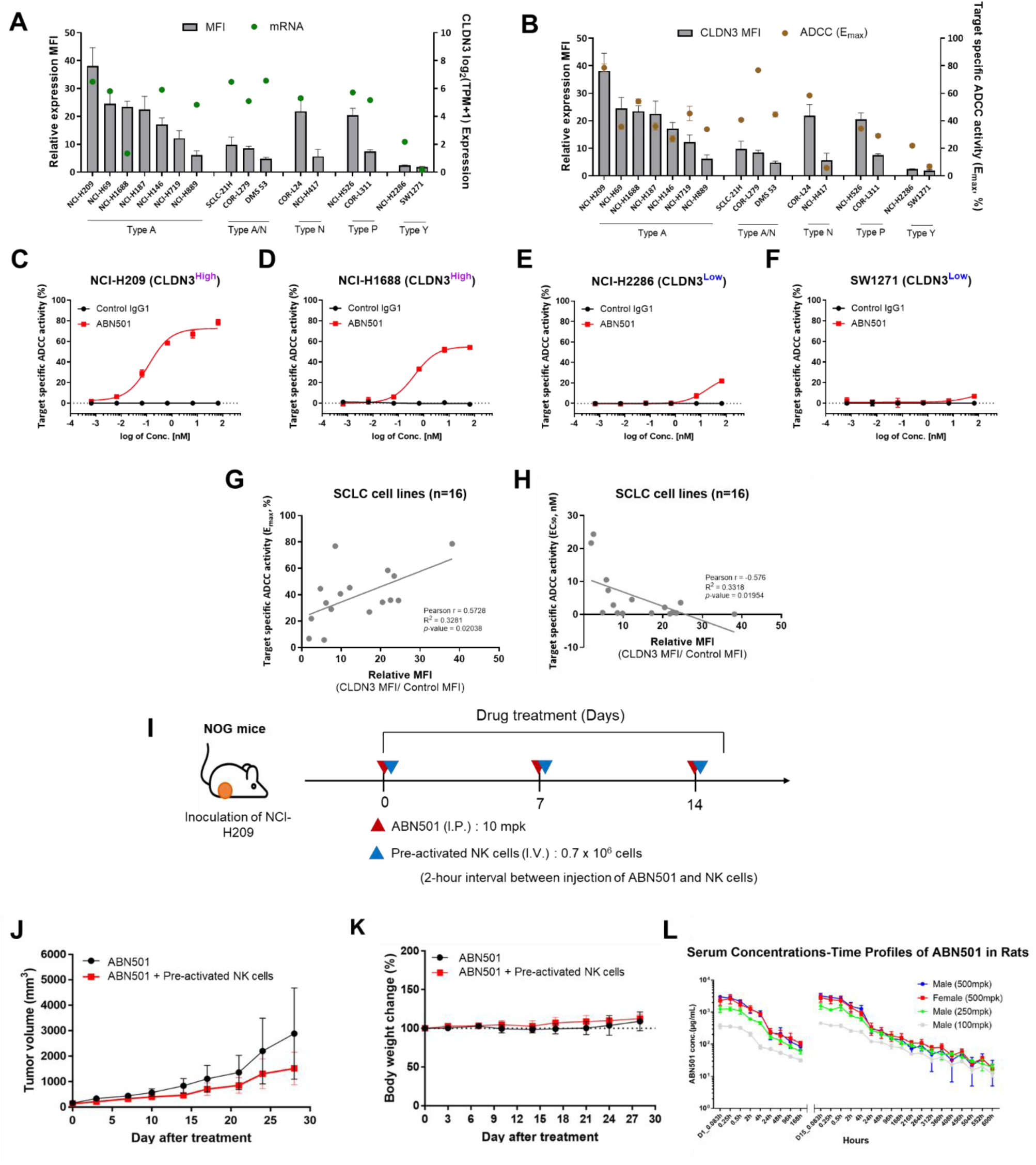
CLDN3-targeted antibody ABN501 induces NK cell–mediated cytotoxicity in vitro and demonstrates antitumor efficacy with favorable safety in vivo. (A) Flow cytometry analysis of CLDN3 surface expression (relative mean fluorescence intensity, MFI; bar graph, left axis) compared with CLDN3 mRNA levels from DepMap (log_2_ TPM+1; blue circles, right axis) across small cell lung cancer (SCLC) cell lines representing different molecular subtypes. (B) Comparison of CLDN3 surface expression (relative MFI; bar graph, left axis) with ADCC activity, expressed as maximum target cell killing (Emax, %; red dots, right axis) following 10 μg/mL ABN501 treatment. (C-F) Representative dose–response curves for cell lines with high CLDN3 expression (NCI-H209, NCI-H1688) or low CLDN3 expression (NCI-H2286, SW1271), incubated for 4 hours with NK-92MI-CD16a effector cells (E:T ratio = 4:1 or 2:1) and ABN501 at indicated concentrations. CLDN3^High^ lines show potent NK-mediated cytotoxicity, whereas CLDN3^Low^ lines demonstrate limited responses. (G–H) Correlation of CLDN3 surface expression with ADCC activity across 16 SCLC cell lines. Higher CLDN3 expression was associated with increased maximal target cell killing (Emax) and lower EC50 values, identifying antigen density as a key determinant of ABN501-mediated NK-cell cytotoxicity. (I) NOG mice were inoculated with NCI-H209 human SCLC cells and treated with intraperitoneal ABN501 (10 mg/kg, once weekly) ± intravenous pre-activated natural killer (NK) cells (0.7 × 10^6^ cells, 2 hours after antibody injection). (J) Tumor volume measurements over 28 days show reduced tumor growth with ABN501 + NK cell co-administration compared to ABN501 alone. (K) Body weight monitoring in NOG mice demonstrates no significant toxicity with either treatment arm. (L) Serum concentration-time profiles of ABN501 in rats (n = 3 per group) following repeated intravenous administration at doses of 100, 250, or 500 mg/kg, showing stable pharmacokinetics across both sexes.

Comparison of CLDN3 expression with NK cell-mediated cytotoxicity showed that surface abundance was the strongest predictor of functional response. Treatment with ABN501, a fully human, afucosylated IgG1 monoclonal antibody engineered to enhance ADCC through NK cell engagement, demonstrated that CLDN3-high cell lines consistently achieved greater maximal cell death (Emax), while low-expressing lines showed limited responses (**Fig. 4B**). Transcriptomic predictions broadly captured CLDN3-high versus CLDN3-low states, and protein-level quantification helped confirm therapeutic susceptibility. Importantly, these findings were consistent across all four SCLC subtypes.

Representative dose-response curves highlighted these findings. CLDN3-high lines such as NCI-H209 and NCI-H1688 demonstrating potent NK cell-mediated cytotoxicity, whereas CLDN3-low lines (NCI-H2286 and SW1271) showed minimal responses (**Fig. 4C-F**). EC50 values further distinguished CLDN3-high from CLDN3-low lines, emphasizing the link between antigen abundance and therapeutic susceptibility. Expanded analyses across individual lines from each subtype confirmed this pattern (**Fig. S5**). Correlation analyses supported this association, with CLDN3 expression positively correlated with maximal target cell killing (Emax; Pearson r = 0.57, p = 0.02) and inversely correlated with EC50 values (Pearson r = −0.58, p = 0.02) (**Fig. 4G–H**), indicating greater sensitivity to ABN501-mediated cytotoxicity in CLDN3-high models. This establishes CLDN3 surface abundance as a key determinant of ADCC efficacy across molecularly diverse SCLC subtypes.

These results demonstrate that CLDN3 is not only consistently enriched in SCLC but also functionally accessible on the tumor surface, where it can be effectively targeted by antibodies such as ABN501 to drive immune effector cytotoxicity. By bridging computational predictions from MadHitter with protein-level and functional validation, these data support CLDN3 as a therapeutically actionable antigen with broad tumor coverage.

### In Vivo Antitumor Efficacy and Preclinical Safety of CLDN3-Directed Therapy

To assess whether CLDN3 targeting translates into therapeutic activity *in vivo*, we tested ABN501 in immunodeficient NOG mice bearing NCI-H209 tumors. Tumor-bearing mice were treated weekly with intraperitoneal ABN501 (10 mg/kg) either alone or in combination with pre-activated NK cells (**Fig. 4I**). ABN501 monotherapy had minimal impact on tumor growth, whereas the combination with NK cells produced marked tumor growth suppression, with tumor volumes remaining essentially flat during the two-week dosing period (**Fig. 4J**). Following cessation of treatment, tumors resumed growth and eventually exceeded 1,000 mm^3^, suggesting that extended dosing could yield more durable control. Body weight remained stable across groups, indicating no overt toxicity (**Fig. 4K**).

To further evaluate pharmacokinetics and safety, ABN501 was administered intravenously to rats at escalating doses (100, 250, or 500 mg/kg) once weekly for three weeks. Serum concentration-time profiling showed predictable, dose-proportional pharmacokinetics across sexes, with stable systemic exposure (**Fig. 4L**). No treatment-related adverse effects were observed: both male and female rats maintained normal growth curves (**Fig. S5 A, B**), consistent food intake (**Fig. S5C, D**), and stable relative organ weights without enlargement or atrophy of critical organs such as liver, heart, kidneys, spleen, and thymus (**Fig. S5E, F**).

Together, these studies demonstrate that CLDN3-directed therapy is well tolerated and capable of driving potent antitumor activity *in vivo*. These findings extend computational predictions, tissue validation, and *in vitro* functional assays into preclinical animal models. By integrating data from transcriptomic prioritization, proteomic profiling of discretely enriched cell populations, cellular cytotoxicity assays, and *in vivo* efficacy and safety, our results establish CLDN3 as a compelling therapeutic target with translational potential in SCLC.

## DISCUSSION

One of the major challenges in treating SCLC has remained inter- and intratumoral heterogeneity with its lack of sufficiently selective cell surface antigens. This study identifies and validates CLDN3 as a promising therapeutic target in SCLC through an integrated pipeline spanning computational modeling, transcriptomic and proteomic validation, functional assays, and *in vivo* testing. Using the MadHitter algorithm applied to single-cell RNA sequencing data, CLDN3 emerged as a top-ranked surface antigen with broad tumor coverage and limited normal tissue expression. Independent validation across IHC, mass spectrometry, and cytotoxicity assays confirmed abundant, accessible expression and functional relevance. Extending these findings, a CLDN3-targeted antibody therapy demonstrated antitumor efficacy *in vivo* with a favorable safety profile, supporting the clinical advancement of CLDN3-targeted therapeutics in SCLC.

CLDN3 is one of 27 distinct claudin genes known in mammals, many of which are dysregulated in human cancer^34^. Claudins are a family of transmembrane proteins that are central to epithelial tight junctions^34^. Additionally, CLDN3, which was originally identified in rats in 1991^35^ and later cloned in humans^36^, is also known to serve as a receptor for *Clostridium perfringens* enterotoxin^37–39^, suggesting an intrinsic capacity for receptor-associated biological activity that distinguishes it from targets such as DLL3.

The therapeutic relevance of claudins is highlighted by the family member CLDN18.2. In Phase III trials, the addition of an anti-CLDN18.2 antibody to chemotherapy significantly improved outcomes in advanced gastric and gastroesophageal cancers, leading to its recent regulatory approval^40,41^. CLDN18.2-directed CAR T-cells have also shown encouraging efficacy and safety in refractory gastrointestinal tumors^42^. Similarly, CLDN6-targeted CAR T-cells combined with an RNA vaccine demonstrated early promise^43^, and interest in CLDN4 and related claudins is growing^44,45^. In our dataset, CLDN4 also ranked highly. While it has been studied preclinically, it has been suggested to have roles beyond epithelial tight junctions, raising concerns over potential on-target, off-tumor toxicities, and therefore translation into clinical trials has been slow ^21,22,46–48^. This leaves CLDN3 as the most immediately actionable candidate in SCLC, and our findings extend the claudin-targeting paradigm to SCLC, nominating CLDN3 as a tumor-selective, broadly expressed, and tractable therapeutic target in this highly lethal disease.

While DLL3 and the recent approval of tarlatamab have provided proof-of-concept for antibody-based immunotherapy, its clinical impact is limited by variable expression and incomplete tumor coverage, particularly in relapsed disease. In addition, several other SCLC targets, including B7-H3 and SEZ6, are under active clinical development but did not meet our predefined selection criteria and were therefore not prioritized in this study. In contrast, CLDN3 was more consistently expressed across malignant populations, spanned all four molecular subtypes, and enriched in tumor versus stromal compartments, suggesting broader prevalence and superior therapeutic potential. Multiple lines of evidence support the accessibility of CLDN3. Membranous expression was confirmed in most tumors, ADCC assays demonstrated that expression levels predict NK cell-mediated killing, and *in vivo* therapy with ABN501 plus NK cells reduced xenograft growth without systemic toxicity. Rodent toxicology studies further showed stable exposure and no overt treatment-related adverse effects, despite ABN501 cross-reactivity with murine CLDN3^49^. The anti-CLDN3 monoclonal antibody ABN501 has previously demonstrated highly selective binding to CLDN3-expressing tumor cells, enabling in vivo visualization of CLDN3-positive tumors^49,50^, tumor-selective accumulation and therapeutic efficacy^51^. Intriguingly, despite robust activity in other tumor contexts, the antitumor efficacy of ABN501 monotherapy was more limited in SCLC, suggesting a context-dependent therapeutic response. This may reflect the unique SCLC tumor microenvironment and reduced dependence on intrinsic antibody-mediated cytotoxicity in SCLC, highlighting a potential limitation of Fc-effector–dependent mechanisms. More potent modalities such as ADC or TCE strategies may be required to achieve durable therapeutic responses in SCLC. Collectively, these findings support CLDN3 as a biologically accessible and therapeutically tractable target, consistent with its emerging relevance within the broader claudin family and its potential for therapeutic application in SCLC.

Although CLDN3 expression and function were validated across multiple datasets and preclinical systems, broader evaluation in additional models, including patient-derived xenografts, will be important to assess durability of response and intertumoral heterogeneity. Toxicity studies in rodents were reassuring but may not fully capture off-tumor effects in humans, particularly in gastrointestinal tissues where baseline CLDN3 expression is present. Finally, the therapeutic potential of targeting CLDN3 may vary by platform, such as bispecific T cell engagers, CAR T-cells, or antibody-drug conjugates, which require further systematic evaluation.

In conclusion, these findings establish CLDN3 as a compelling, tumor-selective, and tractable antigen in SCLC. CLDN3-directed therapies may achieve broader and more consistent coverage than DLL3, particularly in relapsed disease. Future studies should define optimal therapeutic modality, evaluate durability of response, and explore combinations with chemotherapy, ICIs, and DLL3-targeted approaches.

## Supporting information

Supplementary Figures S1 through S6

## ACKNOWLEDGMENTS

The authors gratefully acknowledge the patients and their families for their participation and contributions to this study.

## AUTHOR CONTRIBUTIONS

Conceptualization and study design: B.S., J.C., A.A.S., S.L., A.T.

Data acquisition and investigation: M.N., Y.Z., H.J.S., H.Y., S.H., P.D., D.B., P.S., B.W., N.D.B., R.E.M., D.A., L.B., Z.W.-O., C.M., A.-L.M., R.K., Y.H., A.K.S., A.K., S.Z., A.L.H., T.A.A., N.W.B., T.P.C., R.L., S.M.H., S.H., H.Y.P., J.K., S.H.J., H.J.L., Y.K.S., E.R., S.L., A.T.

Data analysis and interpretation: B.S., J.C., A.A.S., Y.C., B.W., N.W.B., T.P.C., E.R., A.T.

Writing – original draft: B.S., J.C., A.A.S., A.T.

Writing – review and editing: All authors.

Supervision and project administration: H.Y., S.H., Y.K.S., S.L., E.R., A.T. Funding acquisition: S.L., Y.K.S., E.R., A.T.

## DATA AVAILABILITY STATEMENT

The data generated in this study are available from the corresponding author upon reasonable request.

## METHODS

### Ethics Statement

All procedures involving human samples were approved by the Institutional Review Board of the National Cancer Institute (NCT01851395, NCT02146170) and Seoul National University (E2301/001-004) and conducted in accordance with the Declaration of Helsinki. All animal studies were performed under protocols approved by the Institutional Animal Care and Use Committee of Seoul National University. and conducted in compliance with institutional and national guidelines for the care and use of laboratory animals.

### Patients and Sample Collection

Human tumor tissue was obtained through two NIH Clinical Center protocols Rapid Autopsy and Procurement of Cancer Tissue (NCT01851395) and Tissue Procurement and Natural History Study of People With Small Cell Lung Cancer (NCT02146170), which were approved by the NIH Institutional Review Board and Ethics Committee. For NCT02146170, patients with SCLC provided informed consent for tumor biopsies and clinical data collection. For NCT01851395, informed consent was obtained from all patients and their next of kin after the patient’s demise, as per the protocol. Rapid autopsies were performed at the NIH Clinical Center’s Pathology Laboratory Autopsy Suite. Tumor was divided into two parts: one for scRNA-seq and the other formalin-fixed and paraffin-embedded (FFPE). Multiple 4–5-micron serial sections from FFPE blocks were prepared. These sections were used for multiplex immunofluorescence (mIF), immunohistochemistry (IHC), and mass spectrometry-based proteomics. Only specimens showing minimal necrosis and well-preserved morphology were included in downstream analyses.

### Single-cell RNA sequencing

Core biopsies and autopsy specimens intended for scRNA-seq were received in the laboratory immediately after extraction and processed utilizing the Human Tumor Dissociation Kit, (Miltenyi Biotec, cat.no.130-095-929) and the gentleMACS™ Octo Dissociator with Heaters (Miltenyi Biotec, cat.no.130-096-427) using 5 ml of enzyme reaction mix for a duration of 1 hour per manufacturer’s instructions. Samples were then passed through a pluriStrainer® 70 µm (pluriSelect, cat.no.43-50070-51). Samples were spun down at 300g for 5 minutes at 4°C, supernatant was removed, and the samples were resuspended in Invitrogen™ eBioscience™ 1X RBC Lysis Buffer (Invitrogen, cat.no. 50-112-9751) dependent on the amount of tissue per sample, after 5 minutes ice-cold PBS was added to quench the reaction. Samples were centrifuged at 300g for 5 minutes at 4°C and supernatant was removed followed by 2X washing with ice-cold PBS. Dissociated samples were immediately processed by the scRNA-seq protocol using Chromium (10X Genomics) instrument and single-cell 5’ reagent kit.

### Annotating Cell Types

We employed a comprehensive approach to identify cell types in scRNA-seq data, beginning with coarse annotation using both manual and computational methods. Initially, cell types were annotated manually by identifying the top differentially expressed genes for each cluster, following criteria from Chan et al. and the Human Protein Atlas, and validated with the SingleR package using BlueprintEncodeData as a reference^52^. InferCNV was employed to confirm the labeling of tumor and non-tumor clusters^53^. For data processing and cell clustering, we utilized Seurat (v4.0.1, R package)^54^. Cells with fewer than 200 detected genes or more than 20% of UMIs mapping to mitochondrial genes were excluded. Harmony was used for data integration across samples, and gene expression levels were normalized using the ‘NormalizeData’ function with parameters: normalization.method = “LogNormalize” and scale.factor = 10000. Highly variable genes were identified with the ‘FindVariableFeatures’ function using the ‘vst’ method, selecting the top 2,000 genes. Gene expression matrices were scaled and centered using the ‘ScaleData’ function with default settings. Raw count matrices were output for MadHitter analysis and subsequently log2 transformed^18^. Clustering involved principal component analysis (PCA) via the ‘RunPCA’ function, using the first 20 principal components to build a shared nearest neighbor (SNN) graph with the ‘FindNeighbors’ function, and performing clustering with the Louvain algorithm via the ‘FindClusters’ function. Annotations were reviewed manually by examining the top-ranked differentially expressed genes for each cluster using the ‘FindAllMarkers’ function with ‘min.pct = 0.25’. The uniform manifold approximation and projection (UMAP) technique was used to visualize the transcriptional profiles in two-dimensional space.

### Preparing Data for Analysis with MadHitter

Single-cell RNA sequencing generated a dataset of 141,982 cells across 49 tumor samples from 25 patients. To ensure adequate representation of both malignant and non-malignant compartments, samples were required to contain at least 10 tumor and 10 non-tumor cells, yielding 39 qualifying samples for downstream analysis.

For each sample, data were partitioned by cell type so that each (sample, cell type) pair was stored in a separate file, with rows corresponding to genes and columns to individual cells. Each entry represented the log_2_-transformed expression value of a gene in a given cell, with 0 indicating no detectable expression. All non-tumor cells from each sample were merged systematically to create one consolidated file per sample containing tumor cells and another containing non-tumor cells, with consistent gene ordering across files.

Filtering and preprocessing were performed in line with prior approaches. First, any rows containing invalid gene symbols were removed based on Human Gene Nomenclature Committee (HGNC) standards (www.genenames.org). Second, cells with fewer than 10% of genes expressed (>90% zero entries) were excluded. Third, gene aliases were harmonized to current official HGNC symbols to ensure consistency across datasets. Fourth, the dataset was reduced to 2,752 genes encoding proteins expressed on the cell surface, defined previously^19^, with minor modifications. Notably, the 10% gene expression filter was not applied to the Chan et al. dataset, but the other homogenization steps were applied.

Two refinements distinguished this analysis from MadHitter analyses in the original paper^18^. We broadened the search space to include all genes encoding surface proteins, not just receptors, recognizing that established CAR T-cell targets such as CD19 and DLL3 are non-receptors. The 10% expression filter was applied across all genes at the outset, rather than post hoc to the surfaceome subset.

### MadHitter Data Analysis

The MadHitter Algorithm accepts single cell expression data from a tumor cohort and outputs a set of cell surface membrane receptors whose targeting can kill cancer cells most effectively while sparing as many as possible non-tumor cells in the tumor microenvironment^18^. To this end, MadHitter employs two key user-defined parameters, including the upper bound on the proportion of non-tumor cells that may be targeted and the lower bound on the proportion of tumor cells that must be targeted. We ran MadHitter with various settings for the upper bound, ub, and the lower bound, lb, mostly (ub = 0.8, lb = 0.1) and (ub = 0.9, lb = 0.05). We used Gurobi (www.gurobi.com) to solve the instances to optimality. We used the ‘greedy’ mode alternative in the MadHitter to quantify how many cells would be predicted to be killed for each single target and for many pairs of targets, even for samples where the targets being studied did not meet at least one of the ub, lb thresholds.

### Validation dataset

The scRNA-seq dataset of a published study was used for validation^29^. The data were collected by profiling 155,098 single-cell transcriptomes from 22 fresh SCLC clinical samples obtained from 19 patients. This included 54,523 SCLC single-cell transcriptomes. Cells were classified into major cell types such as epithelial, mesenchymal, lymphoid, and myeloid through clustering. Within the epithelial compartment, further clustering identified distinct subtypes of SCLC based on the expression of transcription factors ASCL1, NEUROD1, POU2F3, and YAP1. We used the authors’ filtering and cell type annotations in our analysis.

### Immunohistochemistry

Immunohistochemistry stains were performed on a Leica Bond Max auto-stainer with standard DAB protocol. IHC stains (antibodies list needed) were performed at the National Institutes of Health (NIH), Laboratory of Pathology and at the Molecular Histopathology Laboratory, Frederick National Laboratory, NCI according to instructions of the manufacturers. IHC-stained slides were scanned using the Carl Zeiss AxioScan.Z1 microscope equipped with a Plan-Apochromat 20x NA 0.8 objective. For Cytoplasmic and Membranous markers % positive cells out of total cells in a given segment (based on nuclei scoring) were reported. For membrane markers an H-score was calculated based on the equation: 1 x (% of weakly stained) + 2 x (% of moderately stained) + 3 x (% of strongly stained).

### Mass Spectrometry-based Proteomics

For mass spectrometry (MS)-based proteomics profiling, 15 samples were obtained from 11 unique patients who died of metastatic SCLC and were part of the rapid autopsy protocol NCT01851395. Consecutive 8 μm thin tissue sections were generated using a microtome and mounted onto polyethylene naphthalate (PEN) membrane slides. These sections were stained with H&E to aid in cellular identification. Laser microdissection (LMD) was then performed using the LMD7 system (Leica Microsystems) to selectively isolate tumor and tumor microenvironment regions from each of the 15 tissue specimens. LMD-enriched samples were resuspended in 20 µl of 100 mM triethylammonium bicarbonate with 10% acetonitrile and subjected to pressure-assisted trypsin digestion using pressure cycling technology (PCT) in a Barocycler 2320EXT (Pressure Biosciences, Inc.), following previously described methods^55^. Briefly, samples were incubated at 99 °C for 30 minutes, followed by 50 °C for 10 minutes, with SMART trypsin (ThermoFisher Scientific) added at a ratio of 1 μg per 30 mm^2^ of tissue. The lysis and digestion process involved cycling between 45 kpsi for 50 seconds and atmospheric pressure for 10 seconds over a total of 60 cycles at 50 °C. Peptide digests were quantified using the Pierce BCA Protein Assay Kit (ThermoFisher Scientific). For each sample, 10 µg of digested peptides (n=15 LMD-enriched tumor samples, n=13 LMD-enriched TME samples) were labeled with isobaric tandem mass tags (TMT) according to the manufacturer’s instructions (TMTpro 18-plex Isobaric Label Reagent Set, ThermoFisher Scientific). A pooled reference sample, comprising equal amounts of peptides from each patient, was included in each TMTpro-18 multiplex for normalization. Labeled samples were pooled, cleaned using the EasyPep Maxi MS Sample Prep Kit (ThermoFisher Scientific), and fractionated offline into 36 pooled fractions using basic reversed-phase liquid chromatography (bRPLC). The resulting TMTpro-18 fractions were analyzed by liquid chromatography-tandem MS (LC-MS/MS) using a nanoflow LC system (EASY-nLC 1200, ThermoFisher Scientific) coupled online to a Q Exactive HF-X MS (ThermoFisher Scientific), as previously described^56,57^.

### Flow Cytometric Analysis of CLDN3 Expression

Surface expression of CLDN3 was assessed in patient-derived and established cell lines. Adherent lines were detached with enzyme-free dissociation buffer (Gibco), and suspension lines were washed in DPBS. A total of 2 × 10^5^ cells were incubated with ABN501 (10 µg/mL) or an IgG isotype control (10 µg/mL; Jackson ImmunoResearch) in PBS containing 1% FBS for 1 hour on ice. After washing, cells were stained with FITC-conjugated goat anti-human IgG (1:100; Jackson ImmunoResearch) and analyzed on a BD FACSCalibur. Data were processed with FlowJo, and CLDN3 expression was quantified as mean fluorescence intensity (MFI) relative to isotype control.

### In Vitro ADCC

Adherent targets (2.5 × 10^4^/well) and suspension targets (5 × 10^4^/well) were plated in 96-well plates in RPMI 1640 + 5% FBS. NK-92MI-CD16a cells served as effectors. Targets were treated with ABN501 (≤10 μg/mL) and co-cultured with effectors at 4:1 (adherent) or 2:1 (suspension) effector-to-target ratios for 4 hours at 37°C. Cytotoxicity was measured by LDH release (EZ-LDH kit, DoGenBio). Values were normalized to isotype IgG controls, and dose–response curves and EC₅₀ values were calculated with GraphPad Prism 7.

### NK Cell Purification, Pre-Activation, and Cell Culture

Leukoreduction system (LRS) chambers were obtained from healthy donors via the Seoul Southern Blood Collection Center. Peripheral blood mononuclear cells (PBMCs) were isolated from leukoreduction system (LRS) chambers from healthy donors by Histopaque®-1077 density centrifugation. NK cells were enriched using MACS negative selection (Miltenyi Biotec) and confirmed ≥90% CD56^+^CD3^−^ by flow cytometry. Cells were cultured in SCGM medium (CellGenix) with 10% human serum, penicillin-streptomycin, and L-glutamine. Pre-activation was performed overnight with IL-12 (10 ng/mL), IL-15 (20 ng/mL), and IL-18 (100 ng/mL) (PeproTech). Cells were washed and maintained in RPMI 1640 with IL-15 (5 ng/mL) and IL-2 (46 U/mL). *In vivo* experiments used cells on day 7 post-activation.

### *In Vivo* Efficacy of ABN501 in Xenograft Models

Six-week-old female NOG mice (CIEA, Japan) were subcutaneously inoculated with NCI-H209 cells (5 × 10^6^). When tumors reached ~150 mm^3^, mice were randomized into treatment groups. ABN501 was administered intraperitoneally at 10 mg/kg once weekly for two weeks, either alone or in combination with pre-activated NK cells. For combination treatment, pre-activated NK cells (0.7 × 10^6^ per mouse) were infused intravenously two hours after ABN501 dosing (n = 3 per group). Tumor volumes were measured three times weekly with calipers and calculated as volume (mm^3^) = length × width^2^ × 0.5. Animals were monitored daily for body weight and clinical signs, and tumor growth was followed until volumes exceeded 1,000 mm^3^ or mice met humane endpoint criteria.

### Repeated-Dose Toxicokinetic Study of ABN501 in Rats

Sprague-Dawley rats (6 weeks old; SAMTAKOBIO, Korea) were randomized into toxicity and TK groups. ABN501 (purity 99.61%) was formulated in histidine–methionine–sucrose buffer and given intravenously at 0, 100, 250, or 500 mg/kg once weekly for 3 weeks. Animals were monitored daily for clinical signs, weight, and food intake. At necropsy, liver, heart, kidneys, spleen, and thymus were weighed. TK blood samples were collected via jugular vein at multiple time points up to 600 hours post-dose. Serum ABN501 concentrations were quantified by ELISA using recombinant human CLDN3 as capture antigen.

