## Supplementary Figures S1 through S6 for "Proteomic Profiling Identifies CLDN3 as a Tumor-Selective Therapeutic Target in Small Cell Lung Cancer"

### SUPPLEMENTAL FIGURES

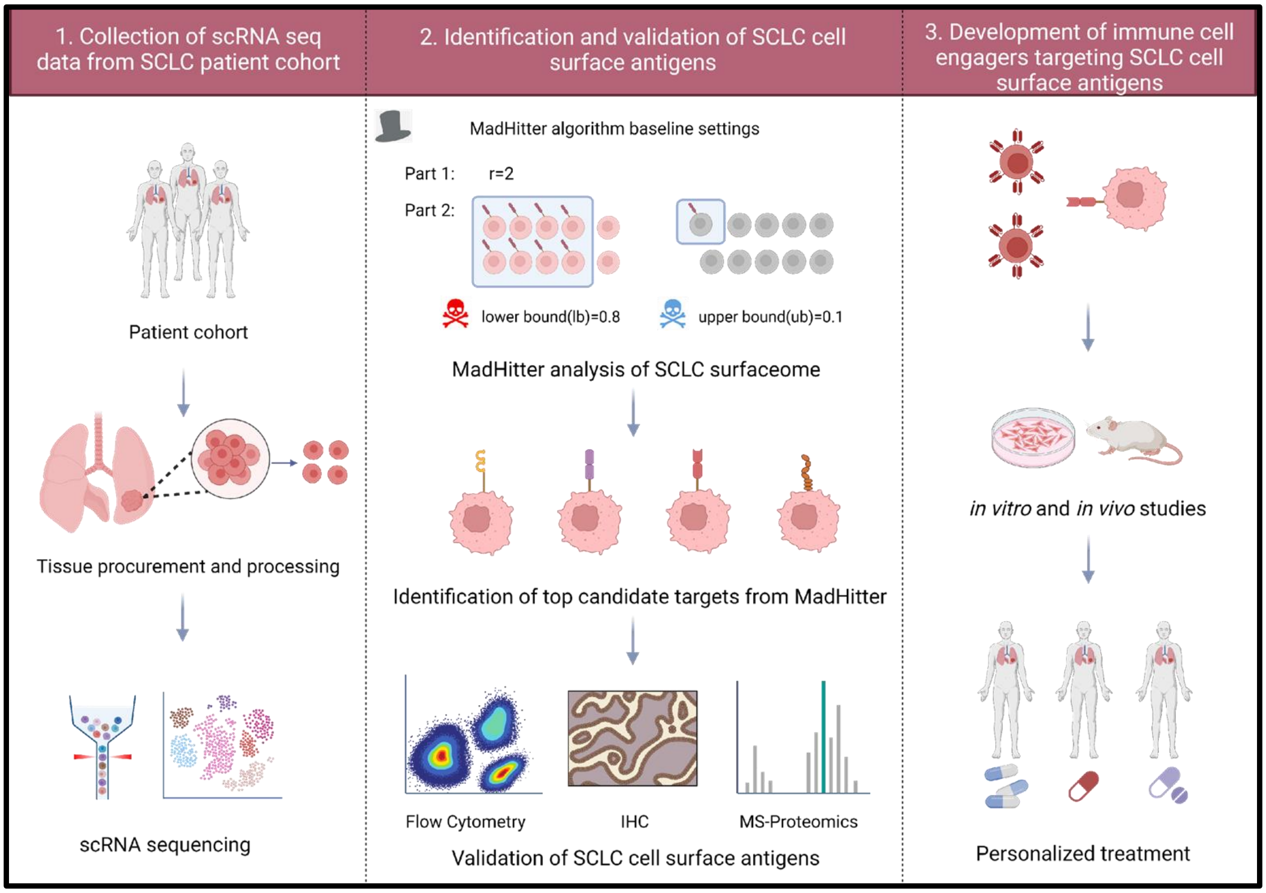


**Figure S1: Schematic overview for targeting SCLC cell surface antigens.** Outline of the workflow for identifying, validating, and developing therapeutic agents targeting cell surface antigens in SCLC using single-cell RNA sequencing (scRNA-seq). **Step 1:** Generation of scRNAseq data from SCLC patients. **Step 2:** Identification and validation of SCLC cell surface antigens: Using the MadHitter Algorithm, which processes scRNA-seq data, the surfaceome of SCLC cells is analyzed. The algorithm works in two stages:
Stage 1: Applies modular baseline settings (e.g., restricting to cell surface proteins) to identify potential therapeutic targets. Stage 2: Focuses on refining these targets by applying upper and lower bounds to balance candidate selection and avoid off-target effects (represented by the hazard symbol). Top candidate targets for SCLC are identified and further validated using immunohistochemistry (IHC) and mass spectrometry (MS)-based proteomics. **Step 3:** Development of antibodies targeting SCLC cell surface antigens. Once validated, the top candidate targets are used to design antibodies that can recognize and bind to the SCLC-specific antigens. These agents are tested through a series of in vitro and in vivo studies to evaluate antigen specificity and antitumor activity. Following successful testing, the validated agents are developed for clinical application in personalized treatment approaches tailored to the specific antigenic profile of each SCLC patient, enhancing therapeutic efficacy while minimizing off-target effects.

| scRNA Sample ID | Cell # | Tissue Type | Collection Type | Site | Specific Site | Race | Gender | Stage at Diagnosis | Prior Txs | Total Txs | Prior Platinum | Prior IO | Platinum Status | PFI | Prior Radiation (Location) |
|---|---|---|---|---|---|---|---|---|---|---|---|---|---|---|---|
| SCAF1355 | 1642 | Tumor | Autopsy | LN | Superior mediastinal | White | M | Limited | 4 | 4 Total | Yes | Yes | Refractory | 0 Days | Yes (Chest |
| SCAF1356 | 2876 | Tumor | Autopsy | Liver | Left Liver lobe mass | White | M | Limited | 4 | 4 Total | Yes | Yes | Refractory | 0 Days | Yes (Chest) |
| SCAF1357 | 708 | Tumor | Autopsy | LN | Superior mediastinal | White | M | Limited | 4 | 4 Total | Yes | Yes | Refractory | 0 Days | Yes (Chest) |
| SCAF1358 | 1951 | Tumor | Autopsy | Liver | Left Liver lobe mass | White | M | Limited | 4 | 4 Total | Yes | Yes | Refractory | 0 Days | Yes (Chest) |
| SCAF1482 | 256 | Tumor | Autopsy | LN | Right Supraclavicular | White | M | Limited | 3 | 3 total | Yes | Yes | Refractory | 0 days | Yes (Chest) |
| SCAF1492 | 794 | Tumor | Autopsy | LN | Right Supraclavicular | White | M | Limited | 3 | 3 total | Yes | Yes | Refractory | 0 days | Yes (Chest) |
| SCAF1814 | 602 | Tumor | Biopsy | LN | Supraclavicular | White | F | Extensive | 1 | 5 total | Yes | Yes | Refractory | 0 Days | No |
| SCAF1815 | 2287 | Tumor | Autopsy | Lung | Right lower lobe lung mas | White | M | Extensive | 2 | 2 Total | Yes | Yes | Resistant | 40 Days | Yes (Brain) |
| SCAF1816 | 1547 | Tumor | Autopsy | LN | Posterior mediastinal | White | M | Extensive | 2 | 2 Total | Yes | Yes | Resistant | 40 Days | Yes (Brain) |
| SCAF1817 | 1809 | Tumor | Autopsy | Liver | Right lobe liver mass | White | M | Extensive | 2 | 2 Total | Yes | Yes | Resistant | 40 Days | Yes (Brain) |
| SCAF1818 | 16698 | Tumor | Autopsy | Adrenal | flank | White | M | Extensive | 2 | 2 Total | Yes | Yes | Resistant | 40 Days | Yes (Brain) |
| SCAF1913 | 3295 | Tumor | Biopsy | LN | Right neck | White | M | Extensive | 4 | 5 Total | Yes | Yes | Resistant | 65 Days | Yes (Brain) |
| SCAF1925 | 2509 | Tumor | Biopsy | Breast | Left breast mass | Multiple | F | Extensive | 3 | 6 total | Yes | Yes | Refractory | 0 Days | Yes (Chest) |
| SCAF2037 | 439 | Tumor | Biopsy | Adrenal | Right adrenal mass | White | M | Limited | 3 | 3 Total | Yes | No | Sensitive | 300 days | Yes (Chest) |
| SCAF2124 | 1544 | Tumor | Biopsy | LN | Supraclavicular | White | M | Extensive | 1 | 2 total | Yes | Yes | Sensitive | 93 days | No |
| SCAF2220 | 7484 | Tumor | Biopsy | Liver | Liver | White | M | Extensive | 2 | 4 total | Yes | Yes | Sensitive | 132 Days | No |
| SCAF2222 | 3633 | Tumor | Biopsy | Liver | Liver | White | M | Extensive | 2 | 4 total | Yes | Yes | Sensitive | 132 Days | No |
| SCAF2225 | 2204 | Tumor | Biopsy | Liver | Liver core biopsy | White | M | Extensive | 2 | 5 total | Yes | Yes | Resistant | 31 Days | Yes (spine) |
| SCAF2229 | 6609 | Tumor | Biopsy | Adrenal | Adrenal gland | White | M | Extensive | 1 | 2 total | Yes | Yes | Sensitive | 174 days | Yes (Brain) |
| SCAF2230 | 2588 | Tumor | Biopsy | Liver | Liver core biopsy | Black | M | Extensive | 2 | 3 Total | Yes | Yes | Resistant | 42 Days | No |
| SCAF2231 | 674 | Tumor | Autopsy | LN | Left Supraclavicular | White | M | Limited | 4 | 4 Total | Yes | Yes | Refractory | 0 Days | Yes (Chest) |
| SCAF2239 | 1232 | Tumor | Autopsy | Lung | Right apex lung mass | White | M | Limited | 3 | 3 total | Yes | Yes | Refractory | 0 days | Yes (Chest) |
| SCAF2326 | 8879 | Tumor | Biopsy | LN | Right cervical | White | M | Extensive | 2 | 3 Total | Yes | Yes | Sensitive | 194 Days | Yes (Chest) |
| SCAF2378 | 5286 | Tumor | Biopsy | Kidney | Kidney mass | Black | M | Extensive | 1 | 2 total | Yes | Yes | Resistant | 53 days | Yes (Brain) |
| SCAF2386 | 13151 | Tumor | Biopsy | Liver | Liver | White | M | Limited | 6 | 6 total | Yes | Yes | Resistant | 20 Days | No |
| SCAF2418 | 3481 | Tumor | Biopsy | Liver | Liver | White | M | Extensive | 4 | 5 total | Yes | Yes | Sensitive | 243 days | Yes (Chest) (Brain) |
| SCAF2419 | 4571 | Tumor | Biopsy | Adrenal | Adrenal gland | White | M | Extensive | 1 | 5 total | Yes | Yes | Resistant | 42 Days | No |
| SCAF2425 | 625 | Tumor | Biopsy | Liver | Liver | Black | F | Extensive | 2 | 3 Total | Yes | Yes | Sensitive | 194 Days | No |
| SCAF2484 | 822 | Tumor | Biopsy | Liver | Liver | White | M | Extensive | 3 | 4 Total | Yes | Yes | Refractory | 0 Days | No |
| SCAF2497 | 10999 | Tumor | Biopsy | LN | Right supraclavicular | White | M | Extensive | 2 | 2 total | Yes | Yes | Sensitive | 93 days | No |
| SCAF2535 | 10979 | Tumor | Biopsy | LN | Right Axillary | White | F | Extensive | 1 | 2 Total | Yes | Yes | Resistant | 62 Days | Yes (Brain) |
| SCAF2538 | 737 | Tumor | Biopsy | LN | Right Axillary | White | F | Extensive | 1 | 2 Total | Yes | Yes | Resistant | 62 Days | Yes (Brain) |
| SCAF2575 | 15144 | Tumor | Biopsy | Liver | Liver | White | M | Extensive | 2 | 6 Total | Yes | Yes | Sensitive | 106 Days | Yes (Chest) |
| SCAF2704 | 4069 | Tumor | Autopsy | Mediastinum | R posterior mediastinal mas | White | M | Extensive | 4 | 4 total | Yes | Yes | Sensitive | 132 Days | No |
| SCAF2705 | 4393 | Normal | Autopsy | Lung | R medial normal | White | M | Extensive | 4 | 4 total | Yes | Yes | Sensitive | 132 Days | No |
| SCAF2707 | 816 | Tumor | Autopsy | Mediastinum | Anterior mediastinal mass | White | M | Extensive | 4 | 4 total | Yes | Yes | Sensitive | 132 Days | No |
| SCAF2708 | 1191 | Tumor | Autopsy | Adrenal | Periadrenal mass | White | M | Extensive | 4 | 4 total | Yes | Yes | Sensitive | 132 Days | No |
| SCAF2747 | 2989 | Tumor | Biopsy | Liver | Liver | White | M | Extensive | 3 | 6 Total | Yes | Yes | Sensitive | 106 Days | Yes (Chest) |
| SCAF2775 | 2401 | Tumor | Biopsy | Liver | Liver | White | M | Extensive | 3 | 5 total | Yes | Yes | Resistant | 42 Days | No |
| SCAF2804 | 2008 | Tumor | Biopsy | Liver | Liver | Black | M | Extensive | 2 | 3 Total | Yes | No | Resistant | 18 Days | No |
| SCAF2815 | 5787 | Tumor | Biopsy | Liver | Liver | White | M | Extensive | 3 | 5 total | Yes | Yes | Resistant | 42 Days | No |
| SCAF2841 | 5514 | Tumor | Biopsy | LN | Suprclavicular | White | M | Extensive | 2 | 3 Total | Yes | Yes | Refractory | 0 Days | Yes (LN) |
| SCAF2854 | 7650 | Tumor | Biopsy | Liver | Liver | White | M | Limited | 1 | 3 Total | Yes | Yes | Sensitive | 119 Days | Yes (Chest) |
| SCAF2915 | 3148 | Tumor | Biopsy | Liver | Liver | White | M | Extensive | 5 | 6 Total | Yes | Yes | Sensitive | 106 Days | Yes (Chest) |
| SCAF2930 | 4440 | Tumor | Biopsy | Pleura | Left pleural mass | Asian | F | Limited | 5 | 5 Total | Yes | Yes | Resistant | 78 Days | Yes (Lung) |
| SCAF2997 | 1404 | Tumor | Biopsy | Liver | Liver | White | M | Limited | 2 | 3 total | Yes | Yes | Sensitive | 119 Days | Yes (Chest) |
| SCAF3061 | 1079 | Tumor | Autopsy | LN | Left hilar mass | White | F | Extensive | 5 | 5 total | Yes | Yes | Refractory | 0 Days | Yes (LN) |
| SCAF3062 | 365 | Tumor | Autopsy | Lung | Left lung peripheral mass | White | F | Extensive | 5 | 5 total | Yes | Yes | Refractory | 0 Days | Yes (LN) |
| SCAF3065 | 3369 | Tumor | Biopsy | Liver | Liver | White | M | Extensive | 6 | 6 Total | Yes | Yes | Sensitive | 106 Days | Yes (Chest) |

**Table S1: Clinical and treatment metadata for SCLC samples analyzed by scRNA-seq.** Metadata for small-cell lung cancer (SCLC) specimens included in this study. Table summarizes clinical and treatment history, including tissue source, anatomical site, stage at diagnosis, prior therapies, platinum sensitivity classification, platinum-free interval, and prior radiation exposure. LN = lymph node, Tx = treatment, IO = immunotherapy, PFI = platinum-free interval.

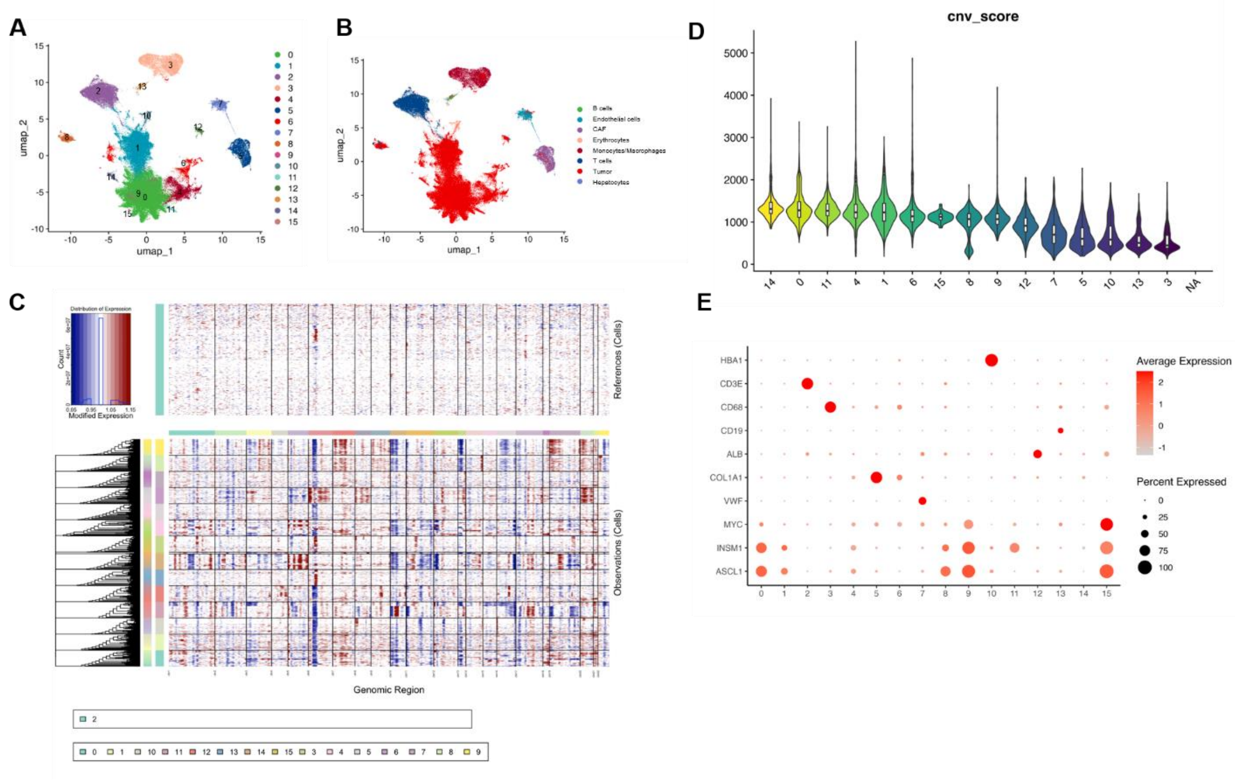


**Figure S2: Identification of malignant cells from scRNA-seq data.** (A) UMAP of all scRNASeq cells (tumor and microenvironment) colored by clusters. (B) UMAP of the epithelial/tumor subset after filtering, showing cluster structure of malignant cells. (C) Heatmap of inferred copy-number variation (CNV) across genomic segments: reference non-malignant cells (top) and tumor cells (bottom); red indicates gains and blue indicates losses. (D) Violin plots of per-cell CNV burden scores across clusters. (E) Dot plot of canonical lineage/marker genes across clusters; dot size reflects the fraction of cells expressing the gene and color reflects average expression.

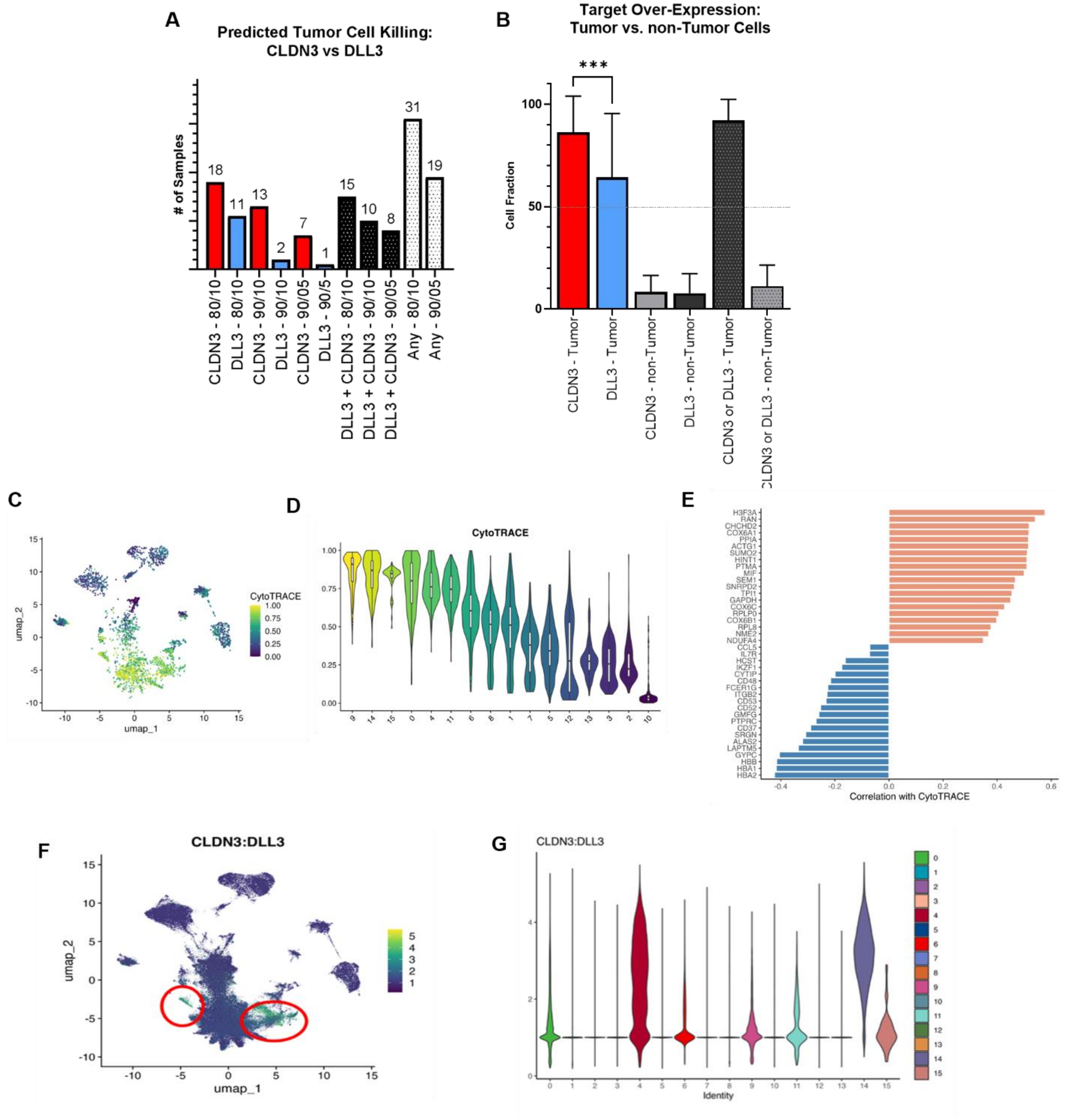


**Figure S3: CLDN3 exhibits broader tumor coverage, lower off-tumor expression, and association with less-differentiated SCLC states.** (A) Predicted tumor cell killing based on combined efficacy (≥80% or ≥90% tumor coverage) and safety (≤10% or ≤5% non-tumor expression) thresholds. CLDN3 satisfied more criteria than DLL3, indicating broader tumor targeting and improved selectivity. (B) Mean % of cells with target overexpression (≥ 0.5) in tumor vs. non-tumor compartments; CLDN3 showed broader tumor coverage with lower non-tumor expression than DLL3. (C) UMAP colored by CytoTRACE scores, indicating relative differentiation state of individual cells (higher scores = less differentiated). (D) Violin plots of CytoTRACE scores by cluster. (E) Bar plot showing correlations between gene-expression programs/ modules and CytoTRACE, highlighting programs associated with less vs. more differentiated states. (F) UMAP of the

CLDN3:DLL3 expression ratio (color scale), marking regions enriched for higher CLDN3 relative to DLL3. (G) Violin plots of the CLDN3:DLL3 ratio across clusters, illustrating heterogeneity of target expression within malignant populations.

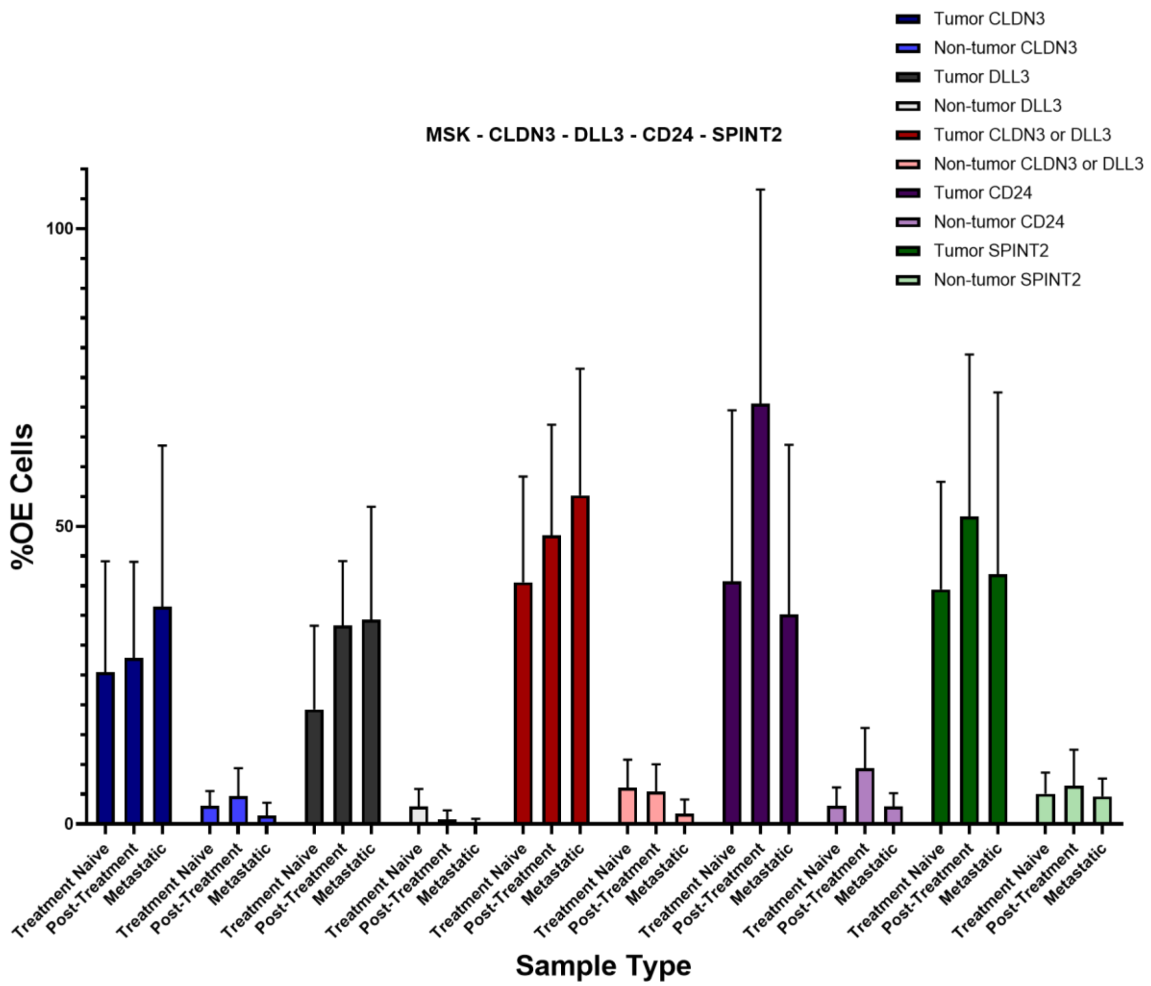


**Figure S4: Tumor and non-tumor cell overexpression of CLDN3, DLL3, CD24, and SPINT2 in the MSK dataset across treatment states.** The proportion of tumor and non-tumor cells overexpressing CLDN3, DLL3, CD24, and SPINT2 in 18 samples from an independent SCLC dataset, categorized into treatment-naïve, post-treatment, and metastatic states. The y-axis represents the percentage of cells overexpressing each target, while the x-axis denotes sample categories. Error bars indicate variability across samples within each group. Tumor cells generally exhibited higher overexpression of CLDN3 and DLL3, with CLDN3 showing greater tumor selectivity. CD24 and SPINT2 were also assessed to explore their expression profiles in tumor versus non-tumor populations.

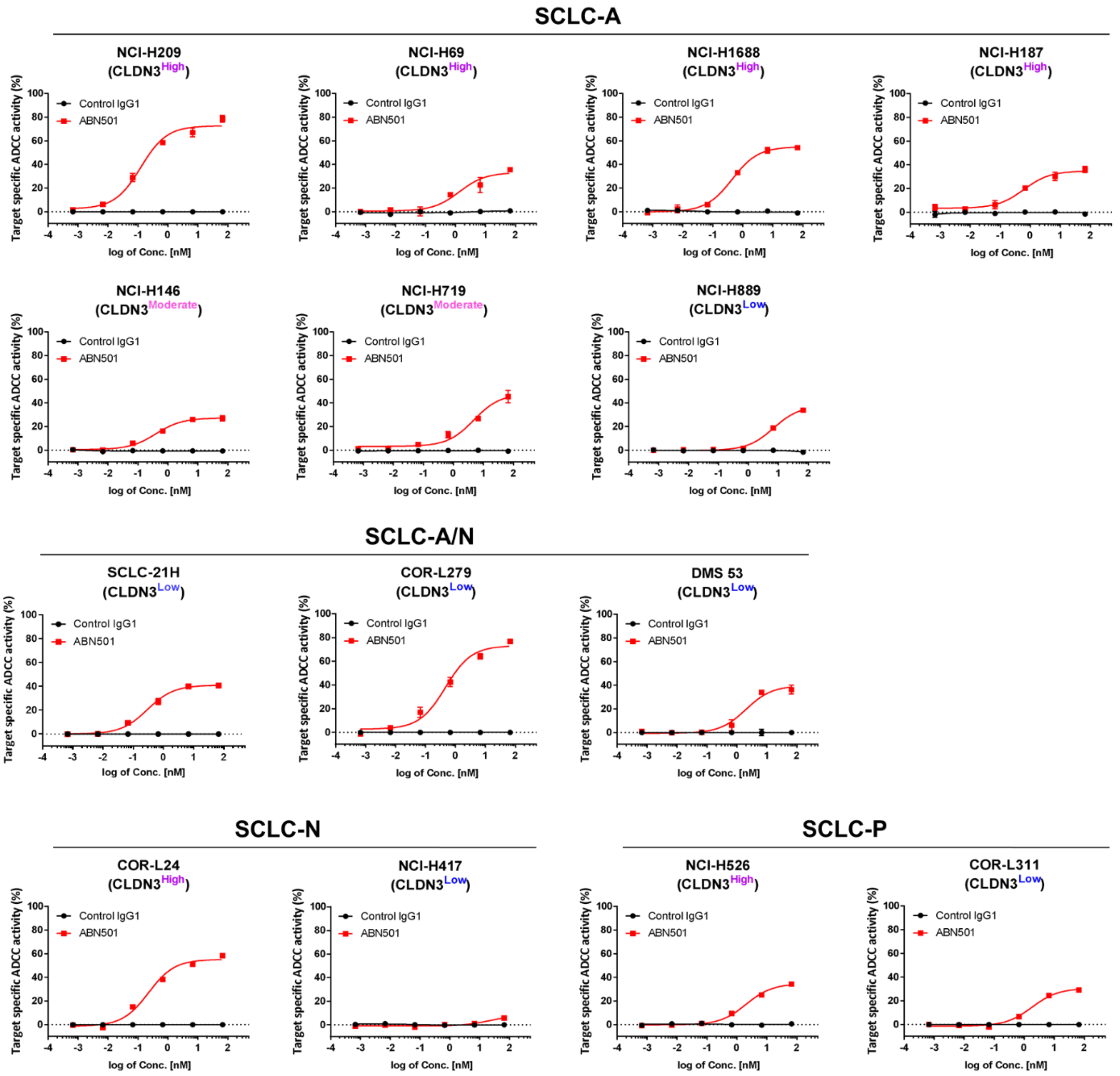


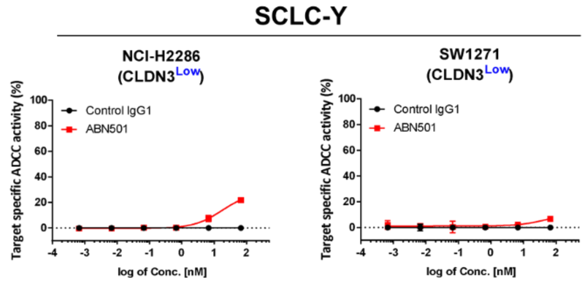


| Cell line | Subtype | Relative MFI | CLDN3 log2(TPM+1) Expression | ADCC (Emax, %) | ADCC ($EC_{50}$, nM) |
|---|---|---|---|---|---|
| NCI-H209 | A | 38.1 ± 6.5 | 6.488 | 78.65 | 0.12 |
| NCI-H69 | A | 24.5 ± 4.0 | 5.815 | 35.70 | 3.59 |
| NCI-H1688 | A | 23.4 ± 2.0 | 1.345 | 54.17 | 0.45 |
| NCI-H187 | A | 22.5 ± 4.7 | N.A | 36.00 | 0.66 |
| NCI-H146 | A | 17.2 ± 2.3 | 5.913 | 26.98 | 0.42 |
| NCI-H719 | A | 12.2 ± 2.7 | N.A | 45.38 | 4.50 |
| NCI-H889 | A | 6.2 ± 1.4 | 4.841 | 33.89 | 7.31 |
| SCLC-21H | A/N | 9.8 ± 2.8 | 6.479 | 40.69 | 0.30 |
| COR-L279 | A/N | 8.5 ± 0.8 | 5.092 | 76.83 | 0.44 |
| DMS 53 | A/N | 4.8 ± 0.6 | 6.561 | 44.64 | 0.53 |
| COR-L24 | N | 21.8 ± 4.1 | 5.314 | 58.42 | 0.23 |
| NCI-H417 | N | 5.7 ± 2.5 | N.A | 5.79 | 10.52 |
| NCI-H526 | P | 20.5 ± 2.4 | 5.724 | 34.37 | 2.14 |
| COR-L311 | P | 7.5 ± 0.6 | 6.561 | 29.11 | 2.85 |
| NCI-H2286 | Y | 2.5 ± 0.2 | 2.176 | 21.96 | 24.36 |
| SW1271 | Y | 1.8 ± 0.3 | 0.151 | 6.76 | 21.68 |

**Figure S5: In vitro CLDN3 NK cell-mediated cytotoxicity across SCLC subtypes.** The SCLC cell lines were grouped by NAPY molecular classification (A, A/N, N, P, Y) and stratified by relative CLDN3 expression (high, intermediate, low). Each cell line was incubated for 4 hours with NK-92MI-

CD16a effector cells (E:T ratio = 4:1 or 2:1) in the presence of ABN501 at indicated concentrations. Dose–response curves show target-specific ADCC activity relative to control IgG. Cell lines with higher CLDN3 expression (e.g., NCI-H209, COR-L24) demonstrated greater sensitivity to ABN501, whereas CLDN3^low lines (e.g., SW1271, NCI-H2286) showed limited cytotoxicity. CLDN3 expression and NK-mediated ADCC across SCLC cell lines, summarized. Values include relative mean fluorescence intensity (MFI), mRNA expression, ADCC Emax (%), and EC50 (nM).

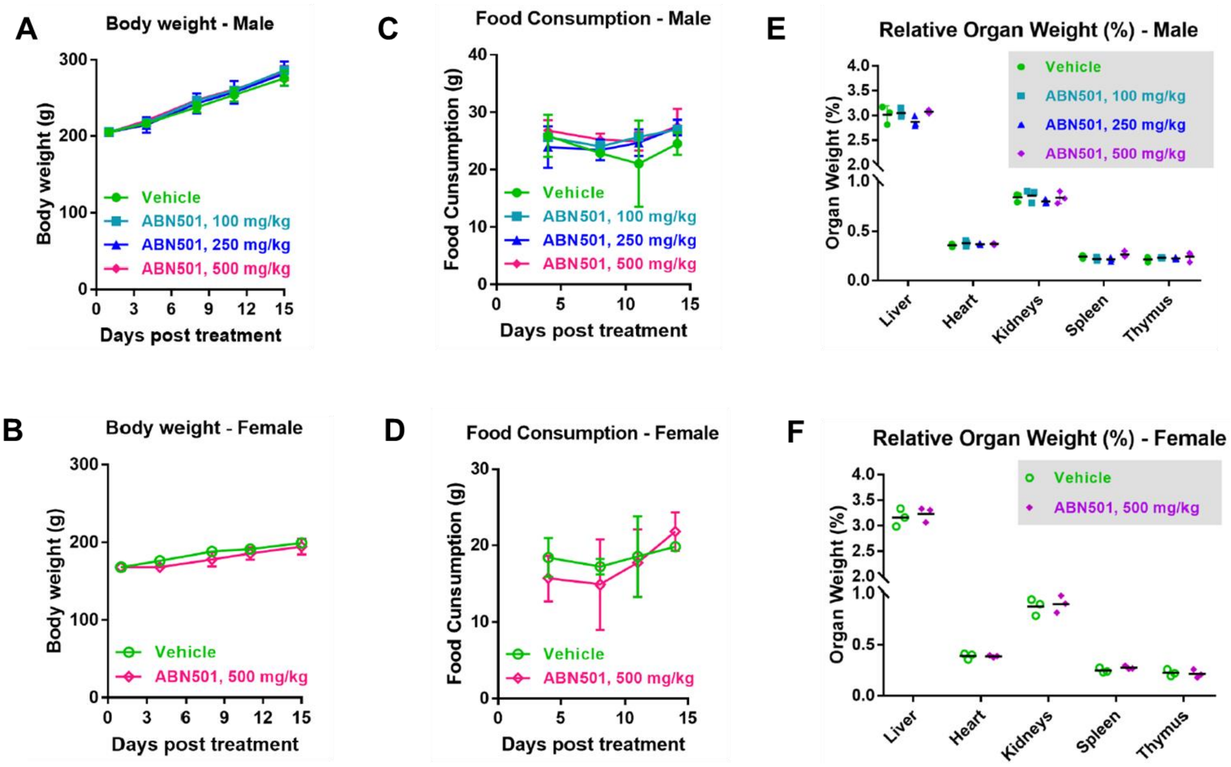


**Figure S6: Effect of ABN501 on body weight, food consumption, or organ weights in male and female rats.** (A-B) Body weight and (C-D) food consumption of male and female rats treated with ABN501 (100-500 mg/kg) weekly for three doses. No weight loss or appetite suppression was observed. (E-F) Relative organ weights at study end showed no enlargement or toxicity-related changes across liver, heart, kidneys, spleen, or thymus in either sex. All dose groups maintained normal growth and food intake over 15 days, indicating no systemic toxicity associated with escalating ABN501 doses.
